# Redox hysteresis controls the NADH-dependent reduction of cytochrome *b*_5_ in rat microsomes

**DOI:** 10.64898/2026.01.08.698363

**Authors:** Oscar H. Martinez-Costa, Ali Ben-Salah, Gabriel N. Valerio, Cristina M. Cordas, Alejandro K. Samhan-Arias

## Abstract

In enzymology, hysteresis is manifested as a time-dependent shift in the kinetic behavior of an enzyme. Through hysteresis, the activation or inhibition of a biological pathway can be regulated by a molecule or metabolite that acts as a hysteretic modulator of the enzyme within that metabolic route. This mechanism of regulation contrasts with those that act on gene expression leading to modulation of enzyme protein levels. Through hysteresis, the amplitude of natural oscillations in metabolic pathways can be adjusted according to the levels of a metabolite that might be beneficial for cells. At physiological level, the slow response of hysteretic enzymes to changes, in the cellular levels of substrates, allows a time-dependent buffering effect on certain metabolites. Understanding the mechanisms and properties of hysteretic enzymes has been important for developing new therapies and improving our understanding of these enzymes in biological systems. However, due to their complex kinetics, the study of hysteretic enzymes has remained a challenge over time.

In this study, we characterized the reduction of cytochrome *b*_5_ by NADH-dependent microsomal enzymes from rat liver using recombinant purified cytochrome *b*_5_, coenzyme Q_10_ and coenzyme Q_0_, as substrates, to mimic the conditions found in biological membranes, where competition between cytochrome *b*_5_ and other substrates might influence their reduction. We found a lag-time-dependent behavior in the cytochrome *b*_5_ reduction compatible with the existence of hysteretic modulation induced by endogenous molecules present in these membranes. Our data suggest that at least for the case of coenzyme Q_10_, fluctuations in its levels may impact metabolic pathways in which reduced cytochrome *b*_5_ levels play a key for the function of the cytochrome *b*_5_-dependent route.

## 1. Introduction

The term hysteresis is defined in physics as the time lag exhibited by a body in responding to external forces. In enzymology this term was early used to describe time-dependent behavior in the kinetic of an enzyme [1]. In general, there are three main molecular mechanisms that appear to be responsible for the manifestation of the hysteresis phenomenon in enzymes, namely, (i) ligand-induced gradual isomerization reaction, (ii) gradual displacement of one allosteric effector by another and (iii) ligand-induced gradual association-dissociation of an enzyme between two forms with differential catalytic activity [2,3]. Hysteresis can regulate the activation or inhibition of a biological mechanism in a manner distinct from those induced by changes in gene expression, by adjusting the amplitude of a metabolic pathway’s natural oscillations, which may be beneficial for the cells [4]. At physiological level, the slow response of hysteretic enzymes to changes in the levels of substances allows a time-dependent buffering effect on certain metabolites within cells. Cells can retain these metabolites for a period before they gradually adapt to changes [5]. This mechanism helps prevent sudden and immediate changes in metabolite concentrations, which is especially important for pathways that share common intermediates or have multiple branching points. As result, the enzyme’s characteristics change over time to match the altered levels of substances in the cell. The buffering capacity of a hysteretic enzyme relies on how quickly it converts one substrate to another. Therefore, hysteretic enzymes are crucial in scenarios where the level of a product or intermediate in a pathway increases rapidly [5]. If changes in substrate concentrations occur more rapidly than the hysteretic response, they may be considered a rapid change. This dynamic is significant, especially when an intermediate is shared between two pathways and undergoes relatively slow changes due to metabolic processes. Responses from hysteretic enzymes can be slow, taking minutes or longer and may lead to substantial changes in the flow of metabolites through a pathway. Understanding the mechanisms and properties of hysteretic enzymes such as pyruvate kinase, phosphofructokinase and choline acetyltransferase has been important for developing new therapies and improving our understanding of these enzymes in biological systems [1,6,7]. However, due to their complex kinetics, the study of hysteretic enzymes has remained a challenge over time.

Ubiquinone, also known as coenzyme Q_10_ (CoQ_10_), is a lipid-soluble compound crucial for cellular energy production. It features a quinone head group and an isoprenoid side chain, rendering it hydrophobic and enabling its integration into lipid membranes. CoQ_10_ is an essential component of the mitochondrial redox chain, shuttling electrons from Complex I to Complex III and establishing the proton gradient necessary for ATP synthesis during cellular respiration. CoQ_10_ is ubiquitous distributed in cell membranes but enriched in organs with high mitochondria content, such as the heart or the liver [8]. A less studied role of this lipophilic molecules, as a regulatory molecule responsible for the hysteretic modulation of mitochondrial Complex I, was early described [9,10]. This study raises the question of whether other cellular NADH-dependent enzymes with the ability to reduce CoQ_10_ might be modulated by hysteresis.

Cytochrome *b*_5_ (C*b*_5_) is a haemoprotein acting as an electron carrier. C*b*_5_ is required for many catalytic reactions including the desaturation and elongation of fatty acids, steroid biosynthesis, drug metabolism and detoxification, as well as oxidative and hydroxylation reactions [11]. NADH-dependent reductive activities of C*b*_5_ and CoQ_10_ exist in different subcellular membranes. In some cases the same protein has been described to be able to catalyze the reduction of both [12], which is consistent with the premise of reactions that can be potentially modulated by hysteresis as occurs in Complex I (two interconvertible states: an active form, rapid, rotenone-sensitive NADH:Q reductase and a slowly activatable form with initial low activity, NEM-insensitive redox centers) which shows biphasic kinetics with a fast initial NADH oxidation followed by slower rates, reflecting a transition from inactive/deactivated to fully active conformation [10].

In this study, we investigated the effect of quinones on the NADH-dependent reduction of C*b*_5_ using microsomes. Our study aims to shed light on the mechanism that regulates this microsomal activity in complex conditions with multiple acceptors and donors.

## 2. Material and Methods

### 2.1 Expression and Purification of Cb_5_

Chemical-competent BL21 E coli strains were transformed with the Heb5 plasmid, containing the codifying sequence for soluble C*b*_5_, and plated on LB agar with 0.1mg/ml ampicillin [13,14]. One colony was picked up and grown for 8 h at 37 °C in 5 mL of LB media supplemented with ampicillin. Then, the culture was transferred into 1 L of Terrific Broth media supplemented by ampicillin and grown for 24 h at 37 °C. After 24 hours, the speed was reduced, and the culture was maintained to be shaken for 12 hours more. Cells were pelleted by centrifugation at 6000 rpm for 30 mins.

For protein purification, pellets were incubated in 50 mM Tris, 1 mM EDTA, 1 mM PMSF pH 8.1, and 0.5 mg/mL lysozyme for 1 h. After lysis, 50 mM of MgCl_2_, 0.2% deoxyribonuclease, 0.5% Triton X-100, 0.5 M NaCl, and 0.05 mg/ml of DNase were added, and the lysate was left under agitation for 1 h. The soluble fraction was separated from debris by centrifugation at 9,000*g* using a JA20 rotor for 30 min at 4 °C. The sample was precipitated with ammonium sulphate up to 50% saturation, was put to agitate for 1 hour, and then centrifuged at 8000*g*. The supernatant was extensively dialyzed against 10 mM Tris, 1 mM EDTA pH 8.1 at 4 °C and loaded onto a diethylaminoethyl (DEAE) Sepharose column (2.5 × 30 cm) (GE Healthcare) previously equilibrated in 10 mM Tris, 1 mM EDTA pH 8.1. A gradient was performed stepwise by the addition of increasing concentrations of Tris buffer (up to 200 mM) to the buffer in the presence of 1 mM EDTA, 0.5% Triton X-100, and 0.2% deoxycholate. C*b*_5_ was eluted in 10 mM Tris, 1 mM EDTA, and 250 mM potassium thiocyanate pH 8.1. Eluent was cleaned from contaminants by the addition of ammonium sulphate 1.1 M and passing it through a CL Sepharose 4B column (2.5 × 10 cm) (equilibrated with 100 mM Tris, 1 mM EDTA) with no retention. After concentration with an Amicon^®^ filter (10-kDa cutoff), the almost pure protein was further separated from contaminants using a Sephadex G75 column (GE Healthcare) (2.5 × 50 cm) equilibrated with Tris 150 mM pH 7.5. After elution, C*b*_5_ was concentrated with an Amicon^®^ filter (10 kDa cutoff) and 30% glycerol was added before freezing at -80 °C.

### 2.2 Preparation of rat microsomes

Three 7-month-old female rats were sacrificed for liver extraction. The livers were minced, washed, and homogenized with a tissue homogenizer (Potter-Elvehjem) in 10 mM Tris buffer, 0.25 M sucrose, 1 mM EDTA, and 1 mM PMSF (phenylmethylsulphonyl fluoride) at pH 8.1. 3 mL of buffer was added for each gram of weighed liver (7.52 g, 9.4 g, and 11.9 g, respectively). After homogenization, the lysate was centrifuged at 960g for 10 minutes at 4°C. The obtained supernatant was centrifuged at 9800g for 20 minutes at 4°C. The second supernatant was centrifuged at 53170g for 60 minutes at 4°C. The microsomal pellet was resuspended in the same buffer as detailed before and the samples were kept at -80 °C until use. The concentration of microsomes was measured using the Bradford method [15].

### 2.3 Preparation of quinones

CoQ_10_ (3 mg/ml) was prepared in 10 mL of ethanol and CoQ_0_ (4 mg/ml) was prepared in potassium phosphate buffer 20 mM pH 7.0. CoQ_10_ and CoQ_0_ spectra were recorded using a Perkin Elmer spectrophotometer in a 1 mL quartz cuvette, filled with 20 mM buffer pH 7.0 and quinones. The extinction coefficient for CoQ_10_ and CoQ_0_ were used to calculate the concentration of these quinones: ε_275nm_ (CoQ_10_) = 14.6 mM^−1^.cm^−1^ [16] and ε_410nm_ (CoQ_0_) = 0.7 mM^−1^.cm^−1^ [17,18].

Supplementation of microsomes with CoQ_10_ was performed by drying the required volume of CoQ_10_ in ethanolic solution and sample resuspension in the presence of microsomes. For membranes +sample manipulation, glass tubes were always used.

### 2.4 Kinetic measurement of Cb_5_ and quinones reduction

C*b*_5_ reduction was measured using a Perkin Elmer Lambda 35 UV spectrometer and a quartz cuvette 10 mm band pass filled with phosphate buffer 20 mM, Diethylenetriaminepentaacetic acid (DTPA) 0.1 mM, pH 7.0, at room temperature in the presence of NADH 150 µM and the concentration of C*b*_5_, quinones, and microsomes indicated in each case. Tracking the absorbance increase at 557 nm. A differential extinction coefficient (reduced-oxidized) for C*b*_5_ of ε_557nm_ = 16.5 mM^−1^.cm^−1^ was used to quantify the amount of reduced C*b*_5_ generated in the cuvette. CoQ_0_ reduction was measured using a Perkin Elmer spectrophotometer in a 1 mL quartz cuvette filled with 20 mM potassium phosphate buffer, 0.1mM DTPA, 1 µg/mL microsomes, 1.2 mM of NADH and different concentrations of CoQ_0_ at 410 nm for 3 min.

### 2.5 Statistics

The data reported in this manuscript are the average of experiments performed by triplicate ± the standard deviation (S.D.).

## 3. Results

### 3.1 UV-vis spectra of electron acceptors of the NADH-dependent microsomal reductases

The UV-vis spectra of CoQ_10_ (50 µM) recorded in aqueous buffer (**Supp. Fig.S1, panel a**) display a maximum absorption band at 282 nm over the absorbance associated with the micellar effect of this compound. CoQ_0_ at the same concentration, shows a maximum band in UV-vis CoQ_0_ spectra at 269 nm, without exhibiting the light scattering effect (**Supp. Fig.S1, Panel b**). A minor band in the UV-vis spectra at 410 nm is observable for CoQ_0_ and is sensitive to the presence of reducers that allows measurement of reductive activities. The visible spectra of C*b*_5_ (10 µM) in the absence and presence of dithionite are shown in **Supp. Fig.S1, panel c** (black and red lines, respectively). Our spectra displayed the characteristic shifts of C*b*_5_ Soret peak and *γ* and *β* bands, induced by reduction, described in the literature [13]. Based on these data, we determined that the ideal wavelength for tracking absorption changes associated with the reduction of C*b*_5_ by microsomes, without interference of CoQ_10_ and CoQ_0_ absorption was 557 nm in case they are added to the assay.

### 3.2 Characterization of the microsomal NADH:CoQ reductase

We selected the band at 410 nm to measure the reduction of CoQ_0_ by microsomes in the presence of NADH (**Fig. 1, panel a)**. The dependence of the microsomal NADH:CoQ_0_ reductase activity (10 µg/mL) on CoQ_0_ concentration followed a hyperbolic trend compatible with a pseudo-first-order kinetics (**Fig. 1, panel b**). The Michaelis-Menten parameters for CoQ_0_ were determined. The average calculated *K_M_* and *V*_max_ values from three different and independent preparations of rat microsomes were 91.3 ± 28.7 µM and the *V*_max_ was 8.5 ± 0.5 µM/min, respectively. Data plotting of the specific activity for the NADH:CoQ_0_ reductase against the microsome concentration was fitted to linear regression that allowed us to obtain a specific activity of 1235 ± 42 µmoles/min/mg of microsomal protein (**Fig. 1, panel c**).

**Figure 1:**
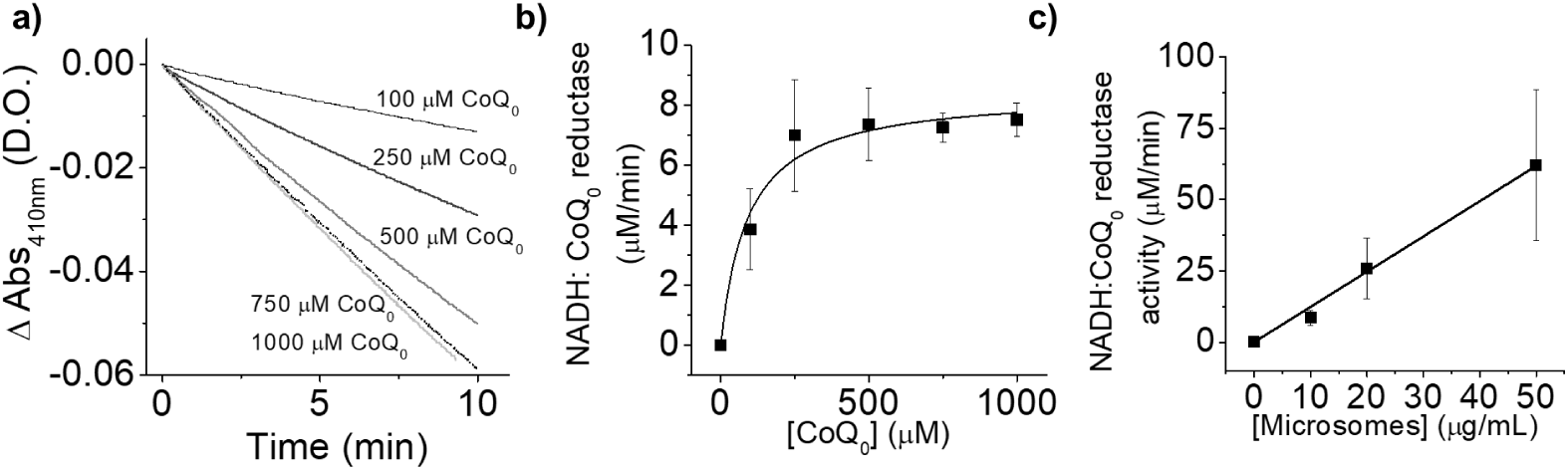
**Panel a: Kinetics of CoQ_0_ reduction by microsomes in the presence of NADH. Panel b: NADH: CoQ_0_ reductase activity of microsomes-dependence on CoQ_0_ concentration:** kinetics experiments were performed in phosphate buffer 20 mM DTPA 0.1mM pH 7.0 in the presence of microsomes 10 µg/mL, NADH 150 µMand increasing concentrations of CoQ_0_ (0-1250 µM). CoQ_0_ reduction was measured by monitoring the absorbance change at 410 nm at room temperature, as described in the Material and Methods section, using a Perkin Elmer Lambda 35 UV Spectrometer and a quartz cuvette of 10 mm bandpass. **Panel c: NADH: CoQ_0_ reductase activity of microsomes-dependence on microsomes concentration:** kinetics experiments were performed in phosphate buffer 20 mM DTPA 0.1mM pH 7.0 in the presence of CoQ_0_ 1mM, a fixed concentration of NADH (150 µM) and increasing concentrations of microsomes using same experimental conditions described in panel b. Experiments shown in this graph are representative experiments of results obtained at least by triplicate ± standard deviation.

### 3.3 Hysteresis mediated by CoQ_0_ on the microsomal NADH-dependent reduction of Cb_5_ is dependent on CoQ_0_ concentration

We measured the kinetics of C*b*_5_ reduction by microsomes in the absence and presence of increasing concentrations of CoQ_10,_ which was incorporated into microsomes, as indicated in the Material and Methods section (**Fig. 2, panel a).** Our experiment showed that the half-life for the reduction of C*b*_5_ (16.7 µM) by microsomes (1 µg/ml) was not affected by the addition of the CoQ_10_ to the membranes. The maximum amount of reduced C*b*_5_ at the stationary stage, in the absence of CoQ_10_ was 5.9 ± 0.4 µM **(Fig. 2, panel a, black line)**. Nevertheless, the maximum concentration of reduced C*b*_5_ reached at the steady state decreased from this value to 5.3 ± 0.4, 3.7 ± 0.7, 2.0 ± 0.2 and 2.4 ± 0.1 in the presence of CoQ_10_ 2, 5, 10 and 15 µM, respectively **(Fig. 2, panel a, red, green, navy blue and cyan lines)**. In addition, we measured the kinetics of C*b*_5_ reduction by microsomes (1 µg/mL) using the water-soluble analog of CoQ_10_, named CoQ_0_ (**Fig 2. panel b)**, as previously indicated in the Material and Methods. The maximum achieved concentration of reduced C*b*_5_ by microsomes (1 µg/mL) in the presence of CoQ_0_ dropped from 5.9 ± 0.4 µM to 3.1 ± 0.4µM, 2.9 ± 0.6 µM, 1.6 ± 0.6 µM and 1.2 ± 0.9 µM in the presence of 2, 5, 10 and 15 µM of CoQ_0,_ respectively (**Fig 2. panel b, red, green, navy blue and cyan lines, respectively)**.

**Figure 2:**
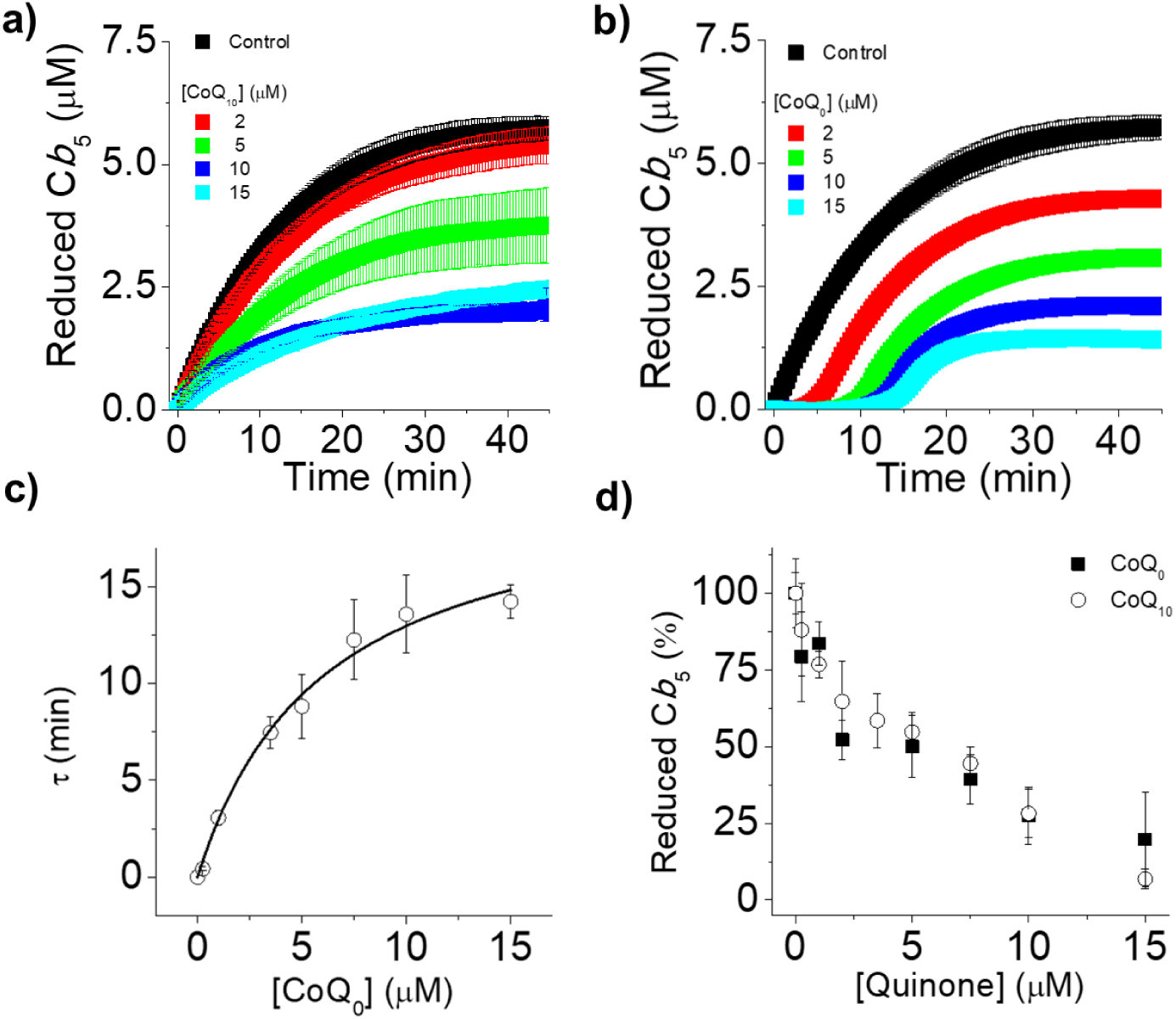
Microsomal C*b*_5_ reduction activity dependence on coenzyme Q concentration. **Panel a**: **kinetics of C*b*_5_ reduction by microsomes in the presence of increasing concentrations of CoQ_10_.** Kinetics experiments were performed using phosphate buffer (20 mM), DTPA (0.1 mM), pH 7.0, in the presence microsomes 1 µg/ml and a fixed concentration of C*b*_5_ (16.7 µM) and NADH 150 µM, at room temperature measuring the difference between reduced and oxidized C*b*_5_ at 557nm, as described at Material and Methods section, using a Perkin Elmer Lambda 35 UV spectrometer and a quartz cuvette 10 mm bandpass. **Panel b**: **Kinetics of C*b*_5_ reduction by microsomes in the presence of increasing concentrations of CoQ_0_**. Kinetics experiments were performed using the same buffer in the presence of microsomes 1 µg/ml and a fixed concentration of C*b*_5_ (16.7 µM) and NADH (150 µM), at room temperature by measuring the difference between the reduced and oxidized C*b*_5_ at 557 nm, as described at material and methods, using a Perkin Elmer Lambda 35 UV Spectrometer and a quartz cuvette 10 mm bandpass. Data shown in panels a and b are representative traces of triplicate experiments. **Panel c: τ value dependence on concentration CoQ_0._** τ values were obtained by calculating the interception of the maximum slope with the x-axis in kinetic experiments in the presence of increasing concentrations of CoQ_0_ Data were fitted to an hyperbolic curve of the type 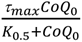 in order to calculate the τ_max_ and the *K*_0.5_ value for CoQ_0_. **Panel d. Dependence of the concentration of C*b*_5_ by microsomes on CoQ concentration (CoQ_10_ and CoQ_0_).** C*b*_5_ reduction activity was determined by measuring the maximum reduced C*b*_5_ concentration in each condition in kinetics experiment. Data shown are mean values ± standard deviation.

In the presence of CoQ_0_, we observed the appearance of initial lag phases in the kinetics of C*b*_5_ reduction by microsomes compatible with the existence of hysteresis, which was dependent on the concentration of CoQ_0_ in the cuvette. Regarding the calculated τ values for each condition, we made a correlation between the concentrations of CoQ_0_ and the τ values obtained in each condition and found a hyperbolic trend with a *K*_0.5_ value of CoQ_0_ of 6.2 ± 1.2 µM of CoQ_0_ and an estimated τ_max_ of 20.0 ± 0.9 min (R^2^= 0.98) (**Fig. 2. panel c, black line**).

We compared the effect of CoQ_10_ and CoQ_0_ on the achieved reduced C*b*_5_ concentration by microsomes (**Panel d, filled black squares and opened black circles, respectively**). Data of reduced C*b*_5_ concentration at different quinone concentrations were normalized to the maximum achieved reduced C*b*_5_ concentration in the absence of quinones. Fitting the inhibition percentage values as a function of quinone concentration to a hyperbolic equation allowed us to calculate the extent of inhibition (**Supp. Fig S2**). The maximum inhibition exerted by both quinones was similar (89 ± 11% and 100 %, for CoQ_0_ by CoQ_10_, respectively), but we found differences in the calculated IC_50_ values: 4.5 ± 1.8 and 2.6 ± 1.1 µM for CoQ_10_ and CoQ_0_, respectively (R^2^= 0.92 and 0.98, respectively). These differences support similar affinities for CoQ_0_ and CoQ_10_ leading to a decrease on the activity of the microsomal system in charge of C*b*_5_ reduction.

### 3.4 Hysteresis on the NADH dependent reduction of Cb_5_ is dependent on the concentration of microsomes

We measured the dependence of the C*b*_5_ reduction rate on microsomal concentration in the presence of C*b*_5_ (16.7 µM) and the sensitivity to 5 µM CoQ_0_ (**Fig 3, panel a).** In the absence of CoQ_0_, the maximum amount of reduced C*b*_5_ dependence on microsomes concentration (1, 2, 10 µg/mL) was 5.9 ± 0.4 µM, 6.9 ± 0.6 µM and 15.2 ± 0.2 µM. In the presence of CoQ_0_ (5 µM), the maximum concentration of reduced C*b*_5_ dropped to 2.9 ± 0.4 µM, 5.4 ± 0.6 µM and 14.4 ± 0.7 µM in experiments performed with the same microsome concentrations, respectively. Kinetic traces were normalized by their respective controls (maximum amount of reduced C*b*_5_ achieved in each condition in the absence of CoQ_0_) and plotted as shown in **Fig 3, panel b**, where 100% reduced Cb5 corresponds to the maximum reduction achieved in each case in the absence of CoQ_0_. The maximum concentration of reduced C*b*_5_ dependence achieved at the stationary stage on microsome concentration in the absence of CoQ_0_ was 36 ± 5 %, 42 ± 4 % and 92 ± 1 % and dropped to 18 ± 3 %, 32 ± 4 % and 87 ± 4 % in the presence of CoQ_0_ (5 µM) to was in experiments performed with 1, 2, 10 µg/mL of microsome, respectively. We plotted the absolute rates dependence for C*b*_5_ reduction (16.7 µM) on microsome concentration, in the absence (black square) and presence of CoQ_0_ (5 µM) (red circle) and found a linear relationship in both conditions with slopes of 0.46 ± 0.02 µM/min and 0.28 ± 0.06 µM/min (**Fig 3, panel c**), which correlates with a 39% inhibition of the activity. These data were normalized to microsomal concentration to obtain the variation of the specific activity with the concentration of microsomes (**Fig 3, panel d**) and a slight increase in the C*b*_5_ reduction rate dependence on microsomes concentration was observed with a slope of 2.0 ± 0.4 nmol/min/mg of protein in absence of CoQ_0_. In the presence of CoQ_0,_ we found a slight decrease in the C*b*_5_ reduction rate dependence on microsomes concentration with a slope of -9.6 ± 4.7 nmol/min/mg of protein, which suggests that CoQ_0_ might interact with microsomal components that enhance the inhibition of the NADH:C*b*_5_ reductase activity induced by CoQ_0_. Finally, initial lag phases from kinetic experiments obtained from Figure 3 were analyzed. We plotted the correlation between τ values with the concentration of microsomes **(panel e)** and fitted the data to an exponential decay model of the type τ = τ_i_*e(-[microsomes]/*k*) + τ_max_, allowing us determination of decay constant value (*τ_1_) of* 9.1 ± 0.9 min and *τ_min_* value of 0.8 ± 0.3 min and a *k_value_ of* 10.6 ± 0.1 µg/mL.

**Figure 3:**
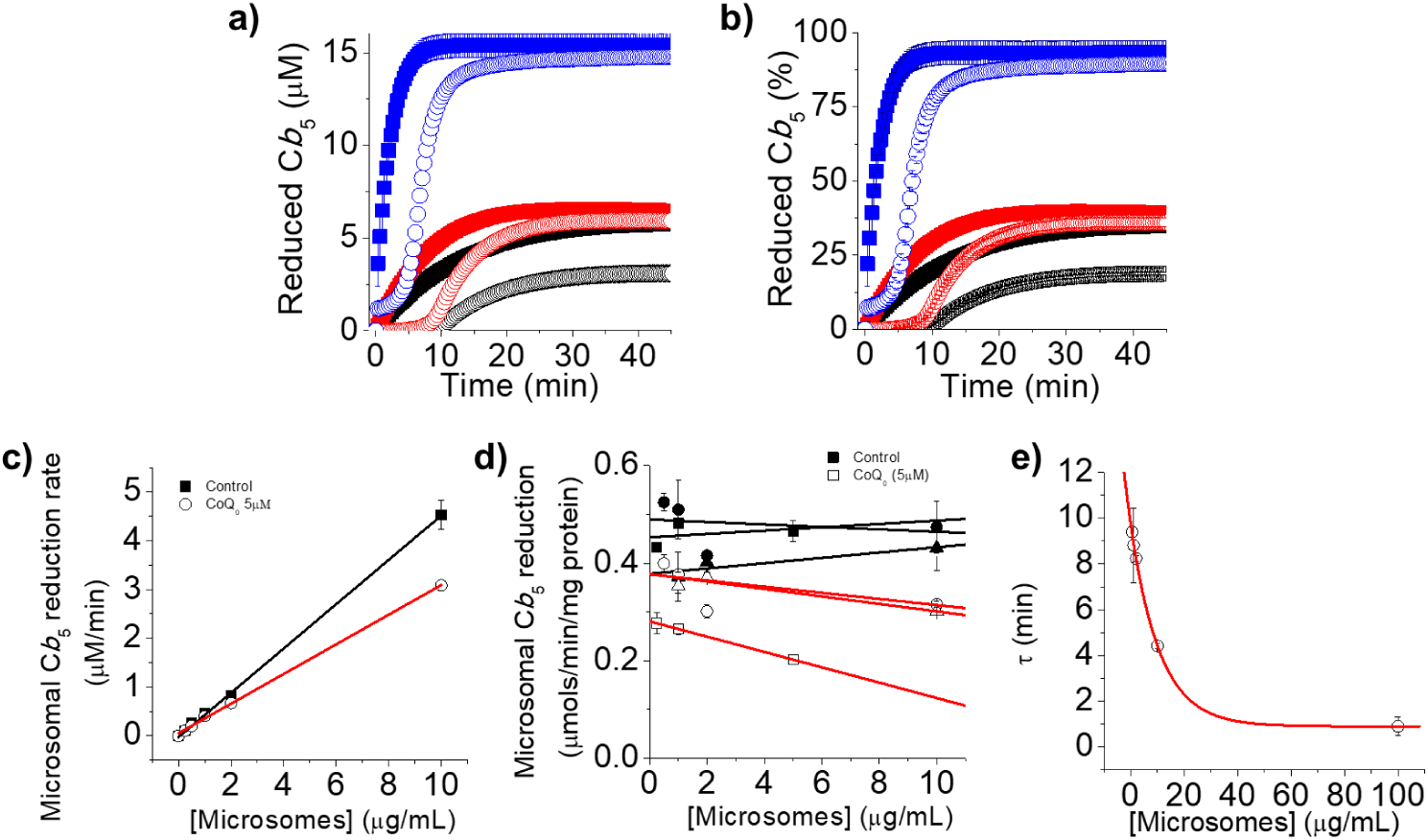
Microsomal C*b*_5_ reduction-dependence on microsomes concentration in the presence of a fixed concentration of CoQ_0_ (5 µM). **Panel a**: **Kinetics of the microsomal C*b*_5_ reduction-dependence on microsomes concentration (1, 2, 10 µg/mL, (black, red and blue symbols, respectively ) in the absence (filled symbols) or presence (open symbols) of CoQ_0_ (5 µM).** Kinetics experiments were performed in phosphate buffer 20 mM, DTPA 0.1 mM, pH 7.0,in the presence of a fixed concentration of C*b*_5_ (16.7 µM) at different concentrations of microsomes 1, 2 and 10 µg/mL (black, red and navy blue colored symbols) and NADH 150 µM, measuring the difference between the reduced and oxidized C*b*_5_, at 557nm, as described in the Material and Methods, using a Perkin Elmer Lambda 35 UV spectrometer and a quartz cuvette 10 mM band pass, at room temperature. Dots represent the average ± SD from representative experiments performed in triplicate using one microsome preparation. **Panel b: Normalized kinetics for the microsomal C*b*_5_ reduction-dependence on microsome concentration in the presence of a fixed concentration of CoQ_0_ (5 µM).** Data from panel a were normalized to the maximum amount of reduced C*b*_5_ achieved in each condition and plotted. Dots represent the average ± SD from representative experiments performed in triplicate using one microsome preparation. **Panel c: Absolute activity for the microsomal C*b*_5_ reduction-dependence on microsomes concentration in the absence (filled squares) and presence of CoQ_0_ (5 µM) (open circles).** The absolute activity was obtained by measuring the maximum slopes of the kinetic traces under each condition. **Panel d: Specific activity for the microsomal C*b*_5_ reduction-dependence on microsomes concentration in the absence (filled squares) and presence of CoQ_0_ (5 µM) (open circles).** The specific activity of microsomal C*b*_5_ reduction was calculated by normalizing the absolute activity to the microsomal concentration used in each assay. Each line represent the linear fitting from experiments with performed in triplicate using one microsome preparation. **Panel e: Dependence of the τ value on the concentration of microsomes in the presence of CoQ_0_ 5µM.** τ values were obtained by calculating the interception of the maximum slope with the x-axis in kinetic experiments where the reduction of C*b*_5_ by microsomes was measured in the presence of increasing concentrations of microsomes in the presence of CoQ_0._ τ values were fitted to the following equation τ = τ_i_*e^(-[microsomes]/*k*)^ + τ_max_ Data shown in panels a, and b are representative traces of triplicate experiments. Data shown are mean values ± standard deviation.

### 3.5 Hysteresis on the NADH dependent reduction of Cb_5_ is dependent on the concentration of Cb_5_

We measured the kinetics of C*b*_5_ reduction by microsomes (1 µg/mL) at increasing concentrations of C*b*_5_ from 0 to 80 µM of C*b*_5_, in the presence of NADH (150 µM). Our data follow an almost linear dependence of microsomal C*b*_5_ reduction activity with the concentration of C*b*_5_ **(Supp Fig. S3)**. Nevertheless, a detailed plotting of the activity at low C*b*_5_ concentrations allowed us to detect the presence of at least two phases consistent with the presence of multiple binding sites for C*b*_5_ in microsomal membranes. The concentration of reduced C*b*_5_ achieved at the steady state at increasing concentrations of C*b*_5_ (3.3, 6.7, 10, 13.4, 16.7, 18.6 and 20.1 µM) was dependent on the used concentration of C*b*_5_ in the assay (black, red, green, navy blue, cyan, pink and olive lines, respectively) (**Fig. 4, panel a**). The maximum achieved C*b*_5_ reduction by microsomes in the presence of CoQ_0_ on the concentration of C*b*_5_ (3.3, 6.7, 10, 13.4, 16.7, 18.6 and 20.1 µM of C*b*_5_) (**Fig. 4, panel b**, black, red, green, navy blue, cyan, pink and olive lines, respectively) with a maximum at 20.1 µM of C*b*_5_ decreased to 67.6 ± 5.8 %. Our data support that in the absence and presence of the quinone the dependence of the maximum achieved concentration of reduced C*b*_5_ at the steady state on C*b*_5_ concentration follows a sigmoidal trend, consistent with the existence of allosteric behavior. We fitted our data to the Hill equation and obtained the following kinetics parameters from 3 independent C*b*_5_ preparations in the absence and presence of CoQ_0_ (5µM) (**Fig. 4, panel c**, black squares and open circles respectively): the estimated maximum achieved concentrations of reduced C*b*_5_ were 9.4 ± 3.0 and 6.5± 1.5 µM, in the absence and presence of CoQ_0_ (5 µM); the estimated concentrations of C*b*_5_ for reaching half of change, we obtained values of 11.4 ± 6.4 and 16.1 ± 3.9 µM in the absence and presence of CoQ_0_.

**Figure 4:**
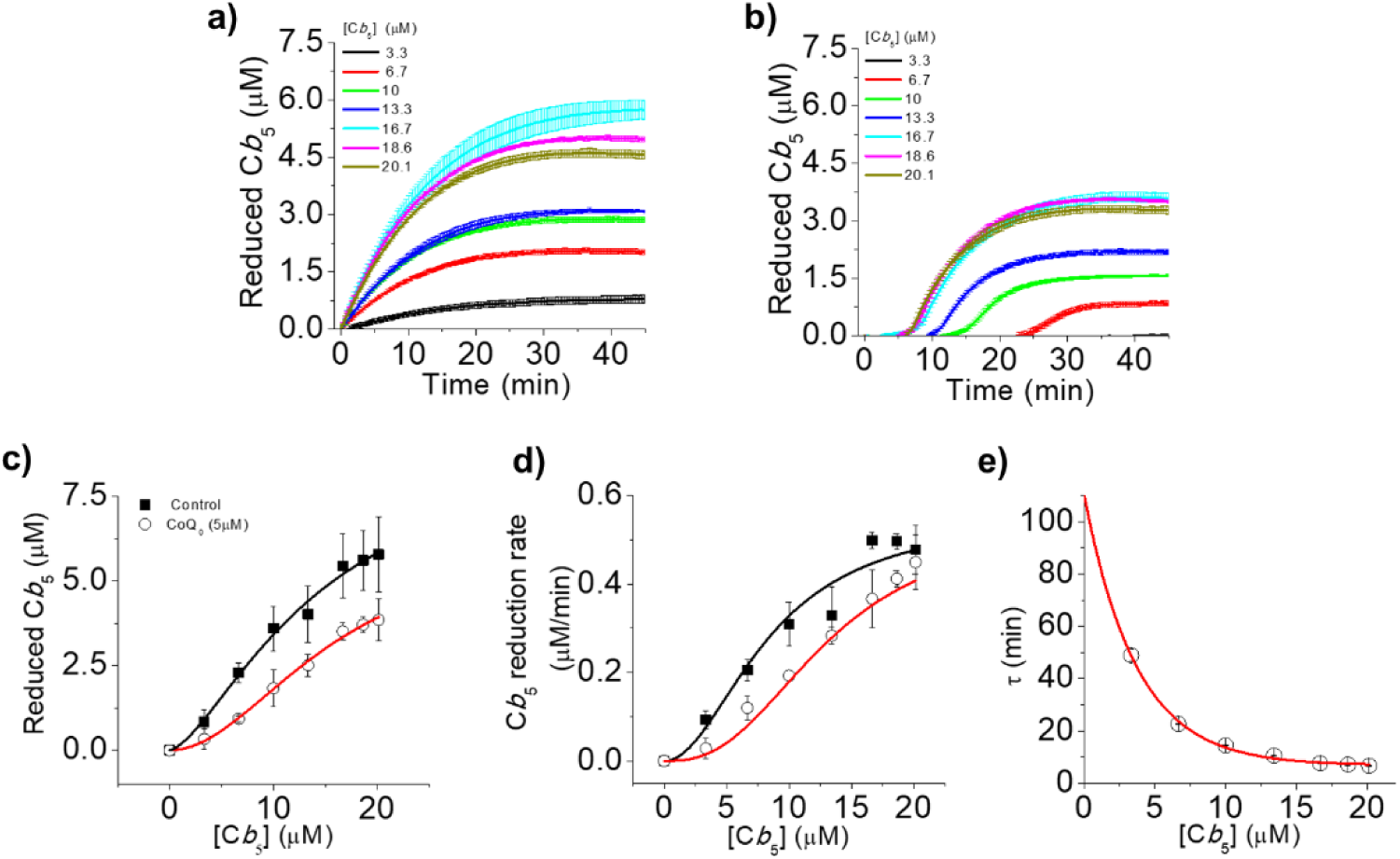
Microsomal C*b*_5_ reduction activity-dependence on C*b*_5_ concentration in the presence of a fixed concentration of quinones (5 µM). **Panel a: Kinetics for the microsomal C*b*_5_ reduction-dependence on concentration of C*b*_5_**. Kinetics experiments were performed in phosphate buffer 20 mM, DTPA 0.1 mM, pH 7.0 in the presence of microsomes 1 µg/ml and increasing concentration of C*b*_5_ 3.3. 6.7, 10, 13.4, 16.7, and 20 µM, and NADH 150 µM, measuring the difference between the reduced and oxidized C*b*_5_ at 557nm, as described in the Material and Methods, using a Perkin Elmer Lambda 35 UV spectrometer and a quartz cuvette 10 mM band pass, at room temperature. Lines represent the average ± SD from representative experiments performed in triplicate using one microsome preparation. **Panel b: Kinetics for the microsomal C*b*_5_ reduction-dependence on concentration of C*b*_5_ in the presence of 5 µM of CoQ_0._** Kinetics experiments were performed using the same conditions indicated for panel a but adding 2 µM of CoQ_0_. Lines represent the average ± SD from representative experiments performed in triplicate using one microsome preparation. **Panel c: Maximum amount of yielded reduced C*b*_5_-dependence on C*b*_5_ concentration in kinetic experiments, in the absence (black open squares) or in the presence of CoQ_0_ (blue open circles).** The maximum amount of reduced C*b*_5_ in each condition was obtained from kinetics experiments performed in panels a and b. **Panel d: Rate of C*b*_5_ reduction activity by microsomes-dependence on C*b*_5_ concentration s, in the absence (black open squares) or in the presence of CoQ_0_ (blue open circles).** C*b*_5_ reduction activity was determined by measuring the maximum rate of C*b*_5_ reduction by microsomes determined by calculating the highest slope in each case. **Panel e: τ value-dependence on concentration C*b*_5_ in kinetic experiments performed in the presence of CoQ_0_ (5 µM).** τ values were obtained by calculating the interception of the maximum slope with the x-axis in kinetic experiments where the reduction of C*b*_5_ by microsomes was measured on the concentration of C*b*_5_, in the presence of increasing concentrations of CoQ_0._ τ values were fitted to the following equation τ = τ_i_*e^(-[C*b*5]/*k*)^ + τ_max_. Data shown in panel a, b, and c are representative traces of triplicate experiments. Data shown are mean values ± standard deviation.

The microsomal C*b*_5_ reduction rate-dependence on C*b*_5_ concentration by microsomes (1 µg/mL), in the absence and presence of CoQ_0_ (5 µM), is shown in **Fig. 4, panel d**. Our data support that both in the absence and presence of CoQ_0_, the dependence of the activity on C*b*_5_ concentration follows a sigmoidal trend, consistent with the existence of allosteric behavior. We simulated our data to the Hill equation and obtained the following kinetics parameters from several batches of purified C*b*_5_: the maximum C*b*_5_ reduction rates-dependence on C*b*_5_ were 0.56 ± 0.04 and 0.54 ± 0.16 µM/min, in the absence and presence of CoQ_0_ (5 µM); *K*_M_ values of 8.4 ± 0.7 and 13.1 ± 1.5 µM, in the absence and presence of CoQ_0_; and *n* values of 2.0 ± 0.4, and 2.6 ± 0.8 µM, in the absence and presence of CoQ_0_. The maximum estimated inhibition percentage in the reduction of C*b*_5_ by microsomes exerted by CoQ_0_ was 31 %. In experiments assayed in the presence of CoQ_0,_ we observed initial lag phases in the kinetic of C*b*_5_ reduction-dependence on C*b*_5_ concentration. We plotted the correlation between τ values and the concentration of C*b*_5_ **(panel e)** and fitted them with an exponential decay equation of the type: τ = τ_i_*e(-[C*b*_5_]/*k*) + τ_max_, where τ_i_ and τ_max_ are the initial and final τ values estimated in the absence of added C*b*_5_ or at saturating C*b*_5_ concentrations and *k* is the concentration of C*b*_5_ at which half of the change in the τ value is achieved. The calculated τ_i_ and τ_max_ values correspond to 110 ± 3 and 5.8 ± 1.0 min for the reduction of C*b*_5_ by microsomes 1 µg/mL and a *k* value of 5.3 ± 1.7 µM, in the presence of CoQ_0_ 5 µM..

### 3.6 Lag phases are present in NADH-dependent reductive events of rat microsomes when a reduction process is measured at 557nm

To assess the biological significance of the findings described in this manuscript we measured the endogenous C*b*_5_ reduction from microsomes by measuring the difference spectra of microsomes (10 µg/mL) and the effect in the absorption at 557 nm after addition NADH. Reduction kinetics are shown in Figure 5. We observed a lag phase preceding the increase in the reduction of a component in microsomes (presumably C*b*_5_ or a cytochrome P450, with absorption at 557 nm), dependent of microsomes concentration. We did not find a significant correlation between calculated τ values and the concentration of microsomes. However, we calculated an average of τ value of 16.9 ± 7 min across different microsomal preparations.

**Figure 5:**
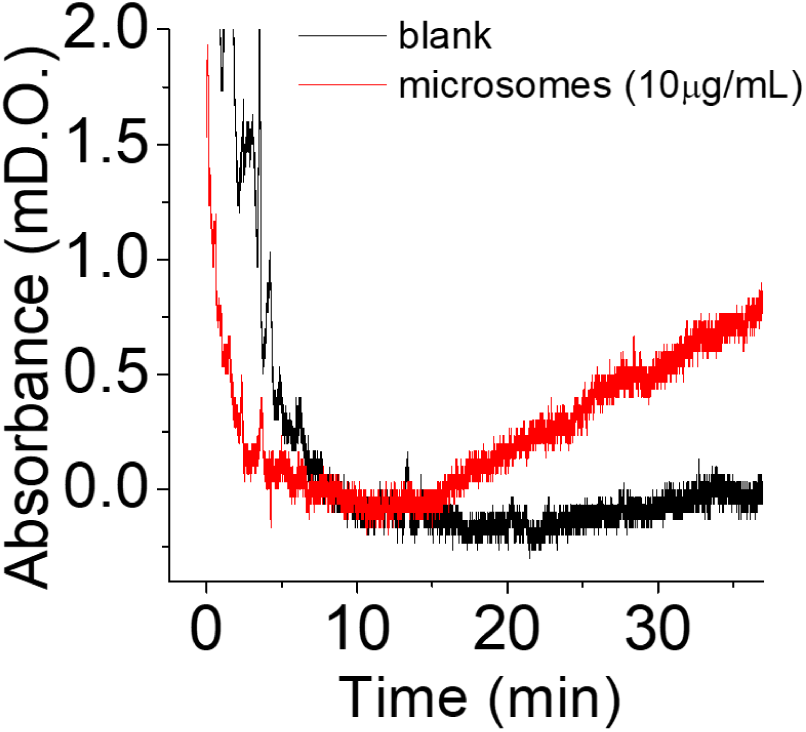
Reduction of endogenous components of microsomes and the presence of hysteresis. **Panel a: Kinetics of microsomal band at 557nm in the presence of NADH and monitored by difference spectra against the same microsomal concentration.** Kinetics experiments were measured by performing a blank of the assay cuvette with phosphate buffer 20 mM DTPA 0.1 mM pH 7.0 in the presence of microsomes 10 µg-mL against the same sample present in the reference cuvette. The assays was initiated by addition of NADH 150 µM to the assay cuvette. A Perkin Elmer Lambda 35 UV Spectrometer was used with a set up wavelength at 557 nm at room temperature and using quartz cuvettes of 10mm bandpass.

These data support that at least a component within microsomal membranes, probably an electron acceptor, exert the hysteretic effect on C*b*_5_ reduction. Most of the characterization performed in this study were performed using relatively high concentrations of C*b*_5_. However, biological C*b*_5_ concentration reported in cells with high amounts of this protein are in the low micromolar range (0.22 µM), much lower than the ones used in our assays. We performed a characterization of the system using a lower concentration of C*b*_5_ (5 µM) keeping the concentration of microsomes (1 µg/mL), NADH (150 µM) and CoQ_10_ and CoQ_0_ (5 µM) constant. The maximum amount of reduced C*b*_5_ after 20 minutes of incubation in the absence of CoQ_10_ was 2.27 µM **(Figure 6, panel a, black line)** and dropped in the presence of CoQ_10_ 0.25, 0.5, 5, and 10 µM, respectively **(Figure 6, panel a, cyan, green, grey and dark grey lines).** We also measured the kinetics of C*b*_5_ reduction by microsomes (1 µg/mL), with a fixed concentration of C*b*_5_ (5 µM), in the presence of increasing concentrations of CoQ_0_ (**Figure 6, panel b**). In the presence of CoQ_0_ (0.01, 0.05, 0.25 and 0.5 µM) the amount of reduced C*b*_5_ decreased (**Figure 6, panel b, red, green, navy and cyan lines, respectively)**. In these experiments, we measured initial lag phases in the presence of CoQ_10_ and CoQ_0_, in a concentration-dependent manner **(Figure 6, panel a and b, lag phases indicated by arrows),** although, lag phases were more pronounced in the presence of CoQ_0_ than in the presence of CoQ_10_. We determined the τ value in the presence of CoQ_0_ and made a correlation with the concentrations of CoQ_0_. The data were fitted to a linear regression and we obtained a slope of 4 min per µM of CoQ_0_ (R^2^= 0.988) (**Figure 6, panel c, red line**). Finally, we compared the effect of CoQ_10_ and CoQ_0_ on the reduction rate of C*b*_5_ by microsomes by normalizing the data to the maximum amount of reduced Cb5 obtained in the absence of quinones (**Figure 6, panel d, filled black squares and red circles, respectively**). The maximum inhibition percentages were 65 ± 3 % 86 ± 1 % for CoQ_10_ and CoQ_0_. The corresponding IC_50_ values were 0.025 ± 0.002 and 0.023 ± 0.014 µM for CoQ_10_ and CoQ_0_, respectively. These data suggest that CoQ_0_ exhibits slightly higher inhibitory affinity than CoQ_10_, as an inhibitor for the microsomal C*b*_5_ reductive system, consistent with the effects observed at higher concentrations of C*b*_5_ (Figure 2).

**Figure 6:**
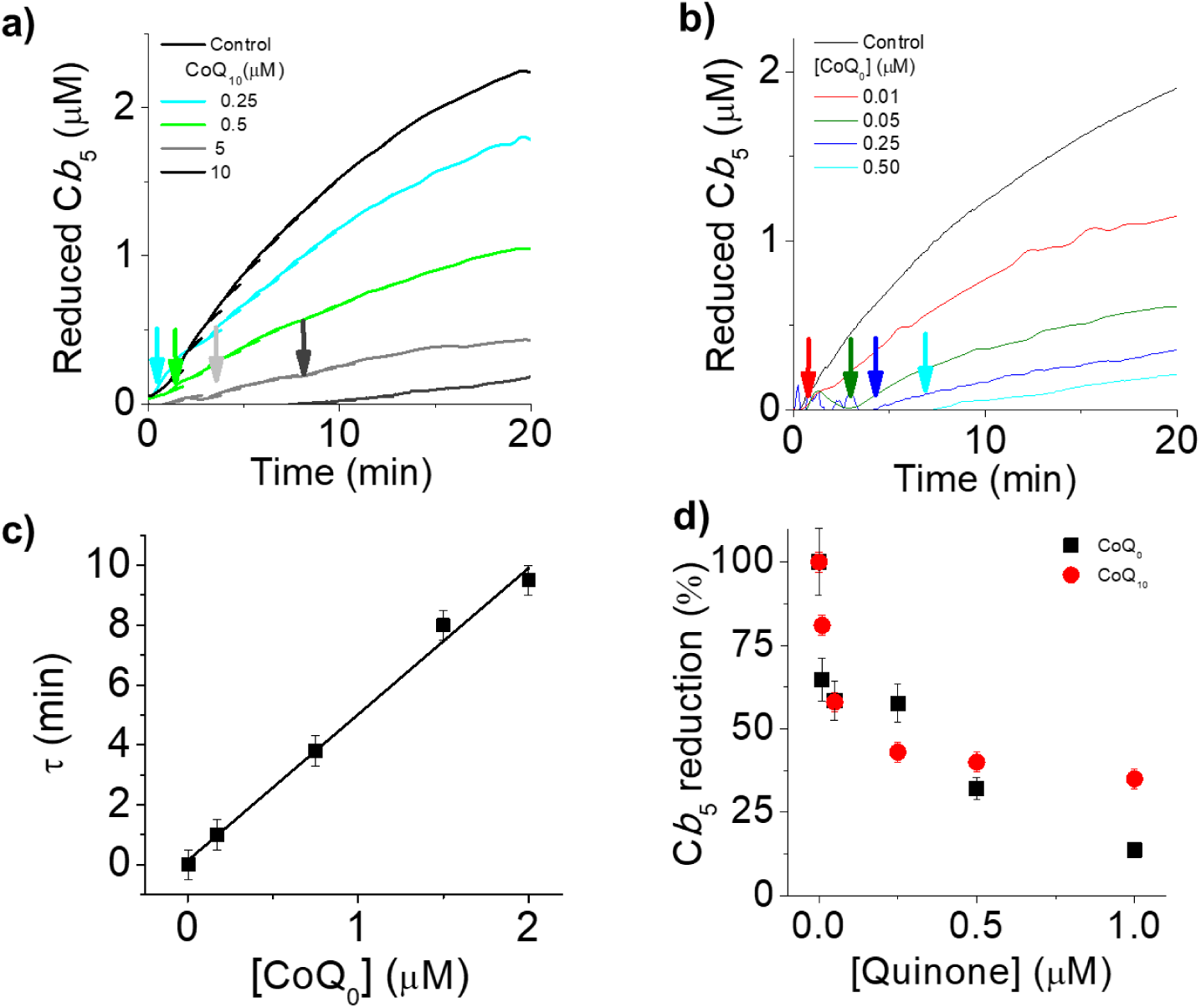
Microsomal C*b*_5_ reduction activity on quinone concentration. **Panel a: Kinetics of C*b*_5_ reduction by microsomes in the presence of increasing concentrations of quinones.** Kinetics experiments were performed using a phosphate buffer 20 mM DTPA 0.1 mM pH 7.0 in the presence of microsomes 1 µg-mL, a fixed C*b*_5_ concentration of 5 µM and NADH 150 µM at room temperature. The increase in absorbance at 557 nm was measured as described at Material and Methods section using a Perkin Elmer Lambda 35 UV Spectrometer and a quartz cuvette of 10mm bandpass. Lines are representative of experiments performed in triplicate. **Panel b: Kinetics of C*b*_5_ reduction by microsomes in the presence of increasing concentrations of CoQ_0._** Kinetics experiments were performed using the same buffer conditions in the presence of microsomes 1 µg/mL and a fixed C*b*_5_ concentration of 5 µM and NADH 150 µM at room temperature. The increase in absorbance increment at 557 nm was measured as described above. Lines are representative of experiments performed in triplicate. **Panel c: τ value-dependence on concentration CoQ_0._** τ values were obtained by calculating the interception of the maximum slope with the x-axis in kinetic experiments where the reduction of C*b*_5_ by microsomes was measured in the presence of increasing concentrations of CoQ_0._ **Panel d: C*b*_5_ reduction activity by microsomes-dependence on coenzyme Q concentration (CoQ_10_ and CoQ_0_, black and red symbols, respectively)** C*b*_5_ reduction activity was determined by measuring the maximum rate achieved in each condition in kinetics experiments shown in panels a and b. Data shown are mean values ± standard deviation and traces shown in panels a and b are representative experiments performed by triplicates.

## 4. Discussion

Membrane NAD(P)H oxidases are surface or membrane embedded proteins that consume cytosolic NAD(P)H for oxidation-reduction reactions occurring in membranes [19]. Many membrane enzymes have adopted the name of electron acceptor used in the reduction reaction they catalyze through the oxidation of NAD(P)H, such as cytochrome P450 reductase (P450R), cytochrome *b*_5_ reductase (C*b*_5_R), quinone oxidoreductase 1 (NQO1) or the thioredoxin reductase . Significant efforts were dedicated to the characterization of these enzymes in which, both biological and artificial electron acceptors were used [20–24]. Some of these compounds may act as electron acceptors enzymes for the same protein with different described enzymatic activities. Nevertheless, and beyond the protein complexity, very little attention has been paid to the use of multiple biological acceptors in the same assay mimicking the physiological environment of biological membranes, as most studies have prioritized the characterization of individual enzymes using a single electron acceptor. In this study using rat liver microsomes, our aim was to characterize the reduction of C*b*_5_ by CoQ_10_ and CoQ_0_, as water soluble analogue of CoQ_10_. A summary of kinetic parameters obtained is shown in Table 1. The major finding of this study is that the C*b*_5_ reduction in these membranes exhibit lag phases, suggesting that the process is hysteresis-dependent. To characterize this phenomenon, we evaluate whether the lag phases, rates of the process and accumulation of reduced C*b*_5_ where dependent on the concentration of substrates (C*b*_5_ and CoQ_0_) and protein concentration, as suggested and reviewed by Neet for the characterization of hysteretic enzymes [25].

We first characterized a NADH:CoQ_0_ reductase activity of microsomes with a *V*_max_ and *K_M_* for CoQ_0_ of 8.5 ± 0.5 µM and 91.3 ± 28.7 µM/min, respectively. Based on the measured *K_M_* value the NADH:CoQ_0_ reductase activity, we can conclude that liver microsomes exhibit low affinity for CoQ_0_ in comparison to other substrates that can be reduced by the addition of NADH. We also characterized the NADH-dependent reduction of C*b*_5,_ which main microsomal contributor to this activity is C*b*_5_R, at 557 nm, a wavelength at which other substrates such as CoQ_0_ does not absorb. The dependence of the microsomal NADH:C*b*_5_ reduction activity on C*b*_5_ concentration follows a linearity and did not reach the activity saturation at tested C*b*_5_ concentrations. We correlated this behavior with presence of enzymes that consume reduced C*b*_5_ in membranes, such as P450s [26], but also with rapid electron transfer between C*b*_5_ molecules, avoiding the saturation of the activity in native membrane systems [27].

Nevertheless, focusing on those data obtained at relative low C*b*_5_ concentrations (1-20µM), we were able to detect sigmoidal trend in the reduction of C*b*_5_, which could be fitted to the Hill equation. We obtained a *K*_M_ value of C*b*_5_ of 8.4 ± 0.7 µM, which falls within the range of described *K_M_* values of C*b*_5_ in the reduction by purified rat C*b*_5_R (11-13 µM [28,29]). A *V*_max_ of 0.56 ± 0.04 µM/min and a cooperativity index of 2 was also determined. An *n* value of 2 suggest the presence of at least two binding sites working in a cooperative manner. As suggested by Neet and collaborators, this type of behavior should be termed hysteretic cooperativity to distinguish it from classical cooperativity arising from site-site interactions [25]. Hysteretic cooperatively is characterized by an enzyme that shifts between states with different catalytic activities and/or by a single-substrate mechanism that, when a second substrate is present at saturating concentrations, effectively behaves as a two-substrate system capable of generating apparent cooperativity [25].

In order to characterize the behavior in the C*b*_5_ reduction by microsomes in the presence of an alternative electron acceptor rather than C*b*_5_, we added CoQ_0_ to our reduction assays. The addition of CoQ_0_ induced the appearance of lag phases in the reduction of C*b*_5_ which is consistent with the existence of a hysteresis process in which CoQ_0_ acts as hysteretic modulator of the microsomal NADH-dependent reduction of C*b*_5_. Increasing concentrations of CoQ_0_ caused a decrease in the achieved C*b*_5_ concentation but also an increase in the τ values, with a maximum of 20.0 ± 0.9 min and a *k* value of CoQ_0_ of 6.2 ± 1.2 min per µM in the presence of C*b*_5_ 16.7 µM.

We also evaluated the dependence of the NADH-dependent reduction of C*b*_5_ on microsomal concentration and determine a specific activity of 0.46 ± 0.02 µmol/min/mg of protein which was inhibited a 39 % in the presence of CoQ_0_ (5 µM). Plotting the specific activity as a function of microsomal concentration revealed the presence of a component in microsomes that decreased the reduction of C*b*_5_ reduction as the concentration of microsomes increase, in the presence of CoQ_0_ (5 µM). We found that increasing the concentration of microsomes induced a decrease of the τ value 0.8 ± 0.3 min with the concentration of microsomes with a *k* value of microsomes for the change of 10.6 ± 0.1 µg/mL, consistent with a hysteretic modulation of the protein responsible for C*b*_5_ reduction. The decrease in the specific activity with increasing microsomal concentration in the presence of CoQ_0_, suggests that there is a protein or non-protein component within microsomes inhibiting this process, possibly by consuming ubiquinol or reduced C*b*_5_. The addition of CoQ_0_ (5 µM) maintained the same *V*_max_ value in the dependence on C*b*_5_ concentration (0.54 ± 0.16 µM/min in respect to 0.56 ± 0.04) but increased the *K_M_* value up to 13.1±1.5 µM, keeping the cooperative value above 2 (n value of 2.6 ± 0.8). Our data might suggest that within the C*b*_5_ concentrations concentration range used, CoQ_0_ acts as a ligand responsible for the observed cooperativity.

We also evaluated whether the hysteretic process observed in the NADH-dependent reduction by microsomes was present in membranes where no additional C*b*_5_ was added to the assays, relying on endogenous haemprotein content. At tested microsomes concentration of 1, 10 and 100µg/mL we observed the presence of lag phases when measured the change of absorbance at 557 nm dependent on the presence of NADH.

Both CoQ_10_ and C*b*_5_ are components of microsomal membranes. In this study we used saturation concentrations of the main substrate C*b*_5_ in order to enzymatically characterize the system. However, the fact that hysteresis was also observed when only endogenous concentrations of C*b*_5_ was present, led us to characterize the hysteresis process using a lower concentrations of C*b*_5_(5µM) than previously used (up to 20µM). Under these conditions, both CoQ_10_ and CoQ_0_ induce hysteresis, correlating with those data reported with higher of C*b*_5_ concentrations.

In membranes, several enzymes display NADH:CoQ_0_ reductase activities, including FSP1 (*K*_M_ 12 µM [30]), NQO1 (0.792 µM [30]), C*b*_5_R (625 µM [31]). Based on the *K*_M_ value for the NADH:CoQ_0_ reductase reported together with the NADH:C*b*_5_ reductive activity described and recent studies with recombinant C*b*_5_R {Citation}, our results suggest that C*b*_5_R is at least one of the enzymes responsible for hysteresis in the reduction of Cb5 by the presence of quinones [32]. Nevertheless, we cannot discard that in complex biological membranes cross reaction between enzymes, including P450R, C*b*_5_R and other enzymes with a NADH:CoQ reductase activity, could be modulated by hysteresis induced by membrane components. Further studies are required to clarify reactions that may be influenced by substrate-induced hysteresis, providing insights into key mechanisms regulating cellular metabolism. In this sense, as described in this study, fluctuations in ubiquinone levels may affect important metabolic pathways, particularly those in which reduction of C*b*_5_ plays an essential functional role.

## Abbreviations

C*b*_5_: Cytochrome *b*₅
CoQ₁₀: Coenzyme Q₁₀
CoQ₀: Coenzyme Q₀
NADH: Reduced nicotinamide adenine dinucleotide
DTPA: Diethylenetriaminepentaacetic acid
EDTA: Ethylenediaminetetraacetic acid
PMSF: Phenylmethylsulfonyl fluoride
DEAE: Diethylaminoethyl

## CRediT authorship contribution statement

**Oscar H. Martínez-Costa:** Writing – review & editing, Writing – original draft, Visualization, Methodology, Investigation, Formal analysis. **Ali Ben-Salah:** Writing , Validation, Supervision, Resources, Methodology, Investigation, Formal analysis. **Gabriel N. Valerio**: Writing – review & editing and conceptualization. **Cristina M. Cordas**: Writing – review & editing and conceptualization. **Alejandro K. Samhan-Arias**: Writing – review & editing, Validation, Supervision, Resources, Project administration, Funding acquisition, and Conceptualization

## Data availability

The data that support the findings of this study are available from the corresponding author, AKSA, on reasonable request.

## Acknowledgement

Open Access funding provided thanks to the CRUE-CSIC agreement with [ACS / Elsevier / IEEE / RSC / Springer / Wiley]. We thank Prof. Carlos Gutiérrez Merino from the Instituto de Biomarcadores de Patologías Moleculares, Universidad de Extremadura, for providing essential material for this work. This work (CMC and GNV) was financed by national funds from FCT - Fundação para a Ciência e a Tecnologia, I.P., under the scope of the project UID/50006/2025 of the Associate Laboratory for Green Chemistry - LAQV REQUIMTE.

---

| | Protein<br>($\mu\text{g/mL}$) | Fixed [$Cb_5$]<br>($\mu\text{M}$) | Fixed [$\text{CoQ}_0$]<br>($\mu\text{M}$) | $K_M$<br>($\mu\text{M}$) | $V_{\max}$<br>($\mu\text{M/min}$) | $n$ value | $\text{IC}_{50}$<br>($\mu\text{M}$) | Activity<br>*umoles/m*<br>*in/mg*<br>*protein* | $\tau$<br>dependence |
| --- | --- | --- | --- | --- | --- | --- | --- | --- | --- |
| NADH:$\text{CoQ}_0$<br>reductase | 1 | - | - | $91.3 \pm 28.7$<br>($\text{CoQ}_0$) | $8.5 \pm 0.5$ | | - | $1235 \pm 42$ | - |
| NADH:$Cb_5$<br>reductase | 1 | 16.7 | - | | | | $2.6 \pm 1.1$<br>($\text{CoQ}_0$)<br>$4.5 \pm 1.8$<br>($\text{CoQ}_{10}$) | | $K_{0.5}(\text{CoQ}_0) =$<br>$6.2 \pm 1.2 \mu\text{M}$<br>$\tau_{\max} =$<br>$20.0 \pm 0.9 \text{ min}$ - |
| NADH:$Cb_5$<br>reductase | 1 | - | 0 | $8.4 \pm 0.7$<br>($Cb_5$) | $0.56 \pm 0.04$ | $2.0 \pm 0.4$ | | | |
| NADH:$Cb_5$<br>reductase | 1 | - | 5 | $13.1 \pm 1.5$<br>($Cb_5$) | $0.54 \pm 0.16$ | $2.6 \pm 0.8$ | | | $\tau_i = 110 \pm 3 \text{ min}$<br>$\tau_{\max} = 5.8 \pm 1.0 \text{ min}$<br>$k(Cb_5) = 5.3 \pm 1.7$<br>$\mu\text{M}$ |
| NADH:$Cb_5$<br>reductase | - | 16.7 | 0 | | | | | $0.46 \pm$<br>$0.02$ | |
| NADH:$Cb_5$<br>reductase | - | 16.7 | 5 | | | | | $0.28 \pm$<br>$0.06$ | $\tau_i = 9.1 \pm 0.9 \text{ min.}$<br>$\tau_{\min} = 0.8 \pm 0.3 \text{ min}$<br>$k(\text{microsomes}) =$<br>$10.6 \pm 0.1 \mu\text{g/mL}$ |

## Notes

### Competing Interest Statement

The authors have declared no competing interest.

## Bibliography

[1] J.L. Rendón, J.P. Pardo, Time-Dependent Kinetic Complexities in Enzyme Assays: A Review, Biomolecules 15 (2025) 641. 10.3390/biom15050641.

[2] J. Madeo, A. Zubair, F. Marianne, A review on the role of quinones in renal disorders, Springerplus 2 (2013) 139. 10.1186/2193-1801-2-139.

[3] C.F. Schaars, A.F.H. Stalenhoef, Effects of ubiquinone (coenzyme Q10) on myopathy in statin users, Curr Opin Lipidol 19 (2008) 553–557. 10.1097/MOL.0b013e3283168ecd.

[4] L. Rinyu, L. Nagy, T. Körtvélyesi, The role of the electronic structure of quinones in the charge stabilization in photosynthetic reaction centers, Journal of Molecular Structure: THEOCHEM 571 (2001) 163–170. 10.1016/S0166-1280(01)00562-0.

[5] C. Frieden, Kinetic Aspects of Regulation of Metabolic Processes: THE HYSTERETIC ENZYME CONCEPT, Journal of Biological Chemistry 245 (1970) 5788–5799. 10.1016/S0021-9258(18)62721-8.

[6] Y. Jiang, X. Li, B.R. Morrow, A. Pothukuchy, J. Gollihar, R. Novak, C.B. Reilly, A.D. Ellington, D.R. Walt, Single-Molecule Mechanistic Study of Enzyme Hysteresis, ACS Cent. Sci. 5 (2019) 1691–1698. 10.1021/acscentsci.9b00718.

[7] C. Frieden, Slow transitions and hysteretic behavior in enzymes, Annu Rev Biochem 48 (1979) 471–489. 10.1146/annurev.bi.48.070179.002351.

[8] V.I. Zozina, S. Covantev, O.A. Goroshko, L.M. Krasnykh, V.G. Kukes, Coenzyme Q10 in Cardiovascular and Metabolic Diseases: Current State of the Problem, Current Cardiology Reviews 14 (2018) 164. 10.2174/1573403X14666180416115428.

[9] A.B. Kotliar, A.D. Vinogradov, [Hysteresis behavior of complex I in delta mu H+-dependent reduction of NAD+ succinate], Biokhimiia 54 (1989) 9–16.

[10] E.O. Maklashina, V.D. Sled’, A.D. Vinogradov, [Hysteresis behavior of complex I from bovine heart mitochondria: kinetic and thermodynamic parameters of retarded reverse transition from the inactive to active state], Biokhimiia 59 (1994) 946–957.

[11] G. Vergéres, L. Waskell, Cytochrome b5, its functions, structure and membrane topology, Biochimie 77 (1995) 604–620. 10.1016/0300-9084(96)88176-4.

[12] D.-H. Hyun, Plasma membrane redox enzymes: new therapeutic targets for neurodegenerative diseases, Arch. Pharm. Res. 42 (2019) 436–445. 10.1007/s12272-019-01147-8.

[13] A.K. Samhan-Arias, L.B. Maia, C.M. Cordas, I. Moura, C. Gutierrez-Merino, J.J.G. Moura, Peroxidase-like activity of cytochrome b5 is triggered upon hemichrome formation in alkaline pH, Biochimica et Biophysica Acta (BBA)- Proteins and Proteomics 1866 (2018) 373–378. 10.1016/j.bbapap.2017.09.010.

[14] A.K. Samhan-Arias, R.M. Almeida, S. Ramos, C.M. Cordas, I. Moura, C. Gutierrez-Merino, J.J.G. Moura, Topography of human cytochrome b5/cytochrome b5 reductase interacting domain and redox alterations upon complex formation, Biochimica et Biophysica Acta (BBA) - Bioenergetics 1859 (2018) 78–87. 10.1016/j.bbabio.2017.10.005.

[15] M.M. Bradford, A rapid and sensitive method for the quantitation of microgram quantities of protein utilizing the principle of protein-dye binding, Anal Biochem 72 (1976) 248–254. 10.1016/0003-2697(76)90527-3.

[16] F.L. Crane, R. Barr, [220] Determination of ubiquinones, in: Methods in Enzymology, Academic Press, 1971: pp. 137–165. 10.1016/S0076-6879(71)18022-6.

[17] I.L. Sun, E.E. Sun, F.L. Crane, D.J. Morré, A. Lindgren, H. Löw, Requirement for coenzyme Q in plasma membrane electron transport., Proceedings of the National Academy of Sciences 89 (1992) 11126–11130. 10.1073/pnas.89.23.11126.

[18] Y. Hatefi, Coenzyme Q (Ubiquinone), in: Advances in Enzymology and Related Areas of Molecular Biology, John Wiley & Sons, Ltd, 1963: pp. 275–328. 10.1002/9780470122709.ch5.

[19] D.J. Morré, A.O. Brightman, NADH oxidase of plasma membranes, J Bioenerg Biomembr 23 (1991) 469–489. 10.1007/BF00771015.

[20] H.W. Strobel, A.V. Hodgson, S. Shen, NADPH Cytochrome P450 Reductase and Its Structural and Functional Domains, in: P.R.O. de Montellano (Ed.), Cytochrome P450: Structure, Mechanism, and Biochemistry, Springer US, Boston, MA, 1995: pp. 225–244. 10.1007/978-1-4757-2391-5_7.

[21] A.K. Samhan-Arias, M.A. Garcia-Bereguiain, F.J. Martin-Romero, C. Gutierrez-Merino, Clustering of plasma membrane-bound cytochrome b5 reductase within ‘lipid raft’ microdomains of the neuronal plasma membrane, Molecular and Cellular Neuroscience 40 (2009) 14–26. 10.1016/j.mcn.2008.08.013.

[22] A.K. Samhan-Arias, C. Gutierrez-Merino, Purified NADH-cytochrome b5 reductase is a novel superoxide anion source inhibited by apocynin: sensitivity to nitric oxide and peroxynitrite, Free Radical Biology and Medicine 73 (2014) 174–189. 10.1016/j.freeradbiomed.2014.04.033.

[23] R.D. Bongard, B.J. Lindemer, G.S. Krenz, M.P. Merker, Preferential utilization of NADPH as the endogenous electron donor for NAD(P)H:quinone oxidoreductase 1 (NQO1) in intact pulmonary arterial endothelial cells, Free Radic Biol Med 46 (2009) 25–32. 10.1016/j.freeradbiomed.2008.09.007.

[24] F. Muller, Chemistry and Biochemistry of Flavoenzymes: Volume III, CRC Press, Boca Raton, 2019. 10.1201/9781351070560.

[25] K.E. Neet, G. Robert Ainslie, [8] Hysteretic enzymes, in: D.L. Purich (Ed.), Methods in Enzymology, Academic Press, 1980: pp. 192–226. 10.1016/S0076-6879(80)64010-5.

[26] C. Bonfils, J.-L. Saldana, C. Balny, P. Maurel, Electron Transfer from Cytochrome b5 to Cytochrome P450, in: E. Arinç, J.B. Schenkman, E. Hodgson (Eds.), Molecular Aspects of Monooxygenases and Bioactivation of Toxic Compounds, Springer US, Boston, MA, 1991: pp. 171–183. 10.1007/978-1-4684-7284-4_10.

[27] D.W. Dixon, X. Hong, S.E. Woehler, A.G. Mauk, B.P. Sishta, Electron-transfer self-exchange kinetics of cytochrome b5, J. Am. Chem. Soc. 112 (1990) 1082–1088. 10.1021/ja00159a030.

[28] M.C. Bewley, C.A. Davis, C.C. Marohnic, D. Taormina, M.J. Barber, The structure of the S127P mutant of cytochrome b5 reductase that causes methemoglobinemia shows the AMP moiety of the flavin occupying the substrate binding site, Biochemistry 42 (2003) 13145–13151. 10.1021/bi034915c.

[29] M.J. Barber, G.B. Quinn, High-level expression in Escherichia coli of the soluble, catalytic domain of rat hepatic cytochrome b5 reductase, Protein Expr Purif 8 (1996) 41–47. 10.1006/prep.1996.0072.

[30] S. Doll, F.P. Freitas, R. Shah, M. Aldrovandi, M.C. da Silva, I. Ingold, A. Goya Grocin, T.N. Xavier da Silva, E. Panzilius, C.H. Scheel, A. Mourão, K. Buday, M. Sato, J. Wanninger, T. Vignane, V. Mohana, M. Rehberg, A. Flatley, A. Schepers, A. Kurz, D. White, M. Sauer, M. Sattler, E.W. Tate, W. Schmitz, A. Schulze, V. O’Donnell, B. Proneth, G.M. Popowicz, D.A. Pratt, J.P.F. Angeli, M. Conrad, FSP1 is a glutathione-independent ferroptosis suppressor, Nature 575 (2019) 693–698. 10.1038/s41586-019-1707-0.

[31] F. Navarro, J.M. Villalba, F.L. Crane, W.C. Mackellar, P. Navas, A phospholipid-dependent NADH-coenzyme Q reductase from liver plasma membrane, Biochem Biophys Res Commun 212 (1995) 138–143. 10.1006/bbrc.1995.1947.

[32] under preparation, Ubiquinone Is a Hysteretic Modulator of the NADH:Cytochrome B5 Reductase Activity of Human Cb5R (n.d.).

