## Supplementary Figures for "Redox hysteresis controls the NADH-dependent reduction of cytochrome *b*_5_ in rat microsomes"

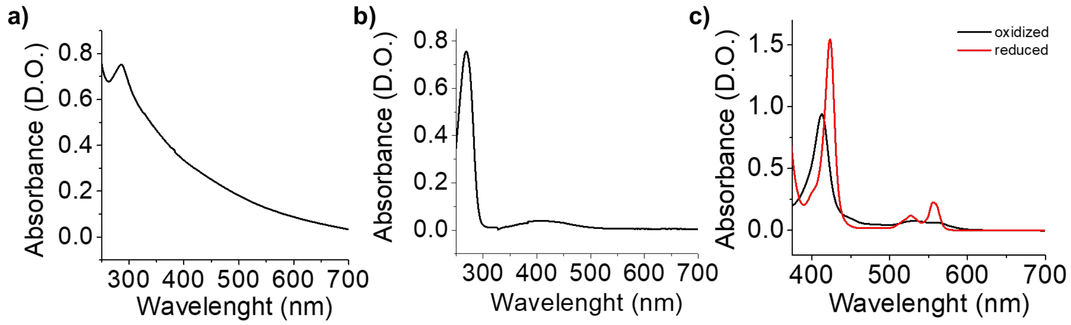


**Supp. Fig.S1: Panel a: $CoQ_{10}$ spectra:** UV-vis spectra of $CoQ_{10}$ were recorded using a Perkin Elmer spectrophotometer in a 1mL quartz cuvette, filled with 20mM potassium phosphate buffer pH 7.0 and 50µM $CoQ_{10}$ at 25ºC. **Panel b: $CoQ_0$ spectra:** UV-vis spectra of $CoQ_0$ were recorded using a Perkin Elmer spectrophotometer in a 1mL quartz cuvette, filled with 20 mM potassium phosphate buffer pH 7.0 and 50µM $CoQ_0$. The wavelength was recorded from 800nm to 200nm. **Panel c:** Oxidized C*b*$_5$ spectra (black line) were recorded using a Perkin Elmer spectrophotometer in a 1mL quartz cuvette, filled with 20 mM potassium phosphate buffer pH 7.0 and C*b*$_5$ 5µM from 800nm to 200nm wavelength. After the measurement of oxidized C*b*$_5$, a few crystals of dithionite were added to the cuvette, and the sample was mixed and the reduced C*b*$_5$ spectra (red line) were measured. Experiments showed in this graph are representative experiments of results obtained at least by triplicate ± standard deviation.

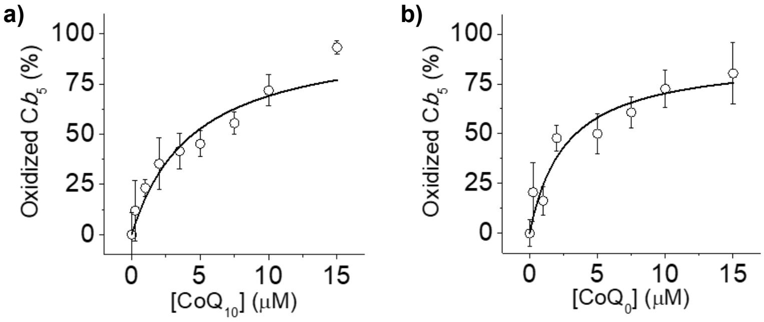


**Supp. Fig S2. Dependence of the oxidized C*b*$_5$ concentration on $CoQ_{10}$ (panel a) and $CoQ_0$ (panel b) after reaching the reduced concentration at the steady state by microsomes.** Panel a and b show the achieved concentration of C*b*$_5$ (%) dependence on the concentration of $CoQ_{10}$ and $CoQ_0$ after data normalization by the maximum amount of reduced C*b*$_5$ achieved in the absence of quinones and after substation of the maximum achieved concentration of reduced C*b*$_5$ (100-percentage of reduced C*b*$_5$)= oxidized C*b*$_5$ concentration. Data were fitted to a hyperbolic curve of the type $\frac{maximum\ percentage\ of oxidized\ [Cb_5][CoQ_0]}{IC_{50}+[CoQ_0]}$ in order to calculate the maximum inhibition percentage and the $IC_{50}$ value for $CoQ_{10}$ and $CoQ_0$. C*b*$_5$ reduction activity was determined by measuring the maximum inhibition percentage in each condition in kinetics experiments shown in panels Figure 2d.Data shown are mean values ± standard deviation.

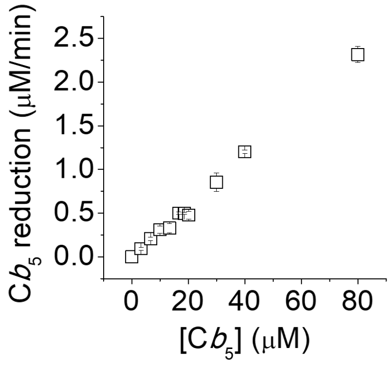


**Supp. Fig.S3: Dependence of the microsomal NADH dependent C*b*$_5$ reduction rate upon the concentration of C*b*$_5$.** Kinetics experiments were performed in phosphate buffer 20 mM, DTPA 0.1 mM, pH 7.0 in the presence of microsomes 1 μg/ml and increasing concentration of C*b*$_5$ 3.3. 6.7, 10, 13.4, 16.7, 20, 30, 40 and 80 μM, and NADH 150 μM, measuring the difference between the reduced and oxidized C*b*$_5$ at 557nm, as described in the Material and Methods, using a Perkin Elmer Lambda 35 UV spectrometer and a quartz cuvette 10 mM band pass, at room temperature.
